# Osmotic Volume Flow in a Donnan System with Permeant Ions

**DOI:** 10.1101/2024.07.22.604593

**Authors:** Gerald S. Manning

## Abstract

Impermeant molecules inside a cell would lead to an inward osmotic flow of water, causing swelling, were it not for the pumping of permeant sodium ions out of the cell as soon as they leak in. The energy barrier model for a semipermeable membrane, first introduced by Debye to provide a molecular-level explanation of the van’t Hoff equation for osmotic pressure, can be used to advantage in this situation, since the pump can be conceptualized as increasing the energy barrier for the sodium ion. The Debye model has previously been extended to include osmosis induced by electrostatically neutral solutes. Discussion of the effect of ion pumping on water transport requires an understanding of osmosis in systems containing permeant ions, that is, Donnan systems. We have obtained an equation for Donnan osmosis across a Debye energy barrier that separates an aqueous solution of permeant sodium, potassium, and chloride ions from a solution containing these permeant ions and additionally an impermeant anion, the latter representing intra-cellular impermeant charged species. Donnan osmosis occurs even if osmolarities on the two sides of the membrane are equal. Numerical representation shows that the Donnan-Debye model provides a quantitative theoretical framework for the action of the sodium/potassium/ATPase ion pump as effectively rendering the extracellular sodium ions impermeant, thus balancing the impermeant molecules inside the cell. Another application of Donnan osmosis shows that ion charge effects, missing from lists of Starling forces, are nonetheless expected to be a major contributor to transport across capillary walls.

**Summary:** Osmosis as driven by Starling forces is applicable only if the solute is electrostatically neutral. For ions, Donnan charge effects dominate. An equation for Donnan osmosis is presented and applied to ion pumps and to transport across capillary walls.

## 1 Introduction

The cells of all organisms contain many molecules dissolved in the cytoplasm that cannot pass through the cell membrane (Milo and Phillips, 2019). They include a diversity of proteins and nucleic acids along with their counterions and also metabolites like ATP and amino acids. These impermeable solutes would give rise to osmotic water flow from the extracellular fluid into the cell, causing swelling and ultimate bursting, unless countered by some mechanism ensuring the stability of the cell. The membranes of plant cells are protected by rigid walls, but the remarkable feature that protects animal cell membranes from osmotic rupture is the action of the Na^+^/K^+^ATPase protein, informally called the sodium pump, embedded in the membrane (Blaustein et al., 2020). In each of its cycles, the pump extrudes 3 Na^+^ions from the cell and takes 2 K^+^ ions inside. The net forced expulsion of Na^+^ ions effectively converts these otherwise permeant ions into impermeant solutes in the extracellular fluid, thus balancing the concentration of impermeant solutes in the cell and preventing osmotic water flow. A primary concern of the present paper is to provide this attractive interpretation (Blaustein et al., 2020) with a quantitative theoretical basis.

The sodium pump will be represented mathematically by implicitly incorporating it into an energy barrier for ion passage across a membrane, just as the set of rate equations known as the “pump-leak” equations (Tosteson and Hoffman, 1960) can use a simple rate term for the pump (Keener and Sneyd, 2009; Aminzare and Kay, 2024). Both rate and energy considerations can provide interesting information on the regulation of cell volume by ion pumping, but a central feature common to both methods of analysis must be a physically consistent mathematical description of the osmotic flow of water induced by a combination of solutes with various degrees of hindered membrane permeability, some or all of them electrostatically charged.

One idea, the physical basis of Starling’s equation for osmotic transport across a biological barrier (Weiss, 1996), is first to take note of van’t Hoff’s criterion, which states that a difference in concentration of an impermeant solute across a semipermeable membrane is balanced at equilibrium by a hydrostatic pressure difference (the “osmotic pressure”). Then, since there is little hydrostatic pressure difference in the biological system of interest, the resulting osmosis, or flow of water volume across the cell membrane, should be proportional to the solute concentration difference. If there is no difference in osmolarity (total concentration of solute particles) there should be no water flow.

The presence of permeant solutes requires modification of this formulation (see Box 3.3 of Blaustein et al., 2020): “With regard to the ability to drive water flow by osmosis, not all solutes are equal. Solutes with low membrane permeability… have far greater osmotic effect than those with high membrane permeability….” It follows that “… two solutions of equal osmolarity can have different osmotic effects on cells.” It can be added that a freely permeant neutral solute contributes nothing to osmosis (by “free” is meant like the solvent water molecules themselves). In the Starling equation, therefore, the concentration of the solute should be multiplied by a “reflection coefficient” ranging from zero for a freely permeant solute to unity for an absolutely inpermeant one (Weiss, 1996). The Starling equation for osmotic volume flow, with reflection coefficients, has a firm thermodynamic basis (Kedem and Katchalsky, 1958), and a mechanistic derivation was found by the present author (Manning, 1968) and reviewed (Manning and Kay, 2023). However, the Starling equation, even with reflection coefficients, is correct only if the solute molecules are electrostatically neutral, for example, like glucose. (For an electroneutral solute like NaCl, which dissociates into its constituent ions, the Starling equation would be correct only if the solute were impermeant and its concentration multiplied by the number of ions in it.)

The Starling equation is not applicable to most physiological systems that include partially permeant ions, where Donnan charge effects are expected to dominate because Coulomb electrostatic forces are both long-range and strong (Donnan, 1911; Scatchard, 1946; Mayer, 1950; Overbeek, 1956; Nagasawa et al., 1959). In the prototype Donnan system, an aqueous salt solution, say NaCl, contains a large protein with net charge, perhaps negative (a net anion). The solution is separated by a membrane from the corresponding salt solution with no protein. The membrane is impermeant to the protein but allows water and the ions of the salt to pass. At equilibrium, there are three Donnan effects, all of them interconnected. The negativeCl^−^ ions are partially excluded from the anionic protein solution while electroneutrality is maintained (Donnan salt exclusion, or ion redistribution). The Donnan electrostatic potential across the membrane is indeed a Nernst potential (Feher, 2018), but only as applied to the specific Donnan ion distribution. When written out in full (see Appendix), it bears little resemblance to the Nernst equation. The Donnan osmotic pressure is influenced by the ion distribution and potential. Again, when written out in full (as in the Appendix), its mathematical representation does not look like the van’t Hoff equation. The Donnan equilibrium effects have their counterparts in the nonequilibrium situation described in this paper. Specifically, the focus here is on what we may call Donnan osmosis to distinguish it from the more familiar van’t Hoff osmosis involving only neutral solutes or impermeant ions.

The model used by Debye (1923) to explain the physical basis of the van’t Hoff equation for equilibrium osmotic pressure represents the semipermeable membrane by a potential energy profile for the solute that rises to infinity as an electrostatically neutral solute molecule approaches the membrane. The infinite amount of energy that must be provided to a solute molecule renders it impermeant. The energy barrier model has been adapted to the nonequilibrium osmotic flow of water induced by neutral solute molecules of intermediate degrees of membrane permeability, up to absolutely impermeant, when a counterbalancing external pressure difference is not allowed to build up (Manning, 1968; Manning and Kay, 2023). It was demonstrated that the water flow is driven by an abrupt pressure drop at the membrane-solution interface which causes a gradient of pressure inside the membrane. This paper presents an analysis of Donnan osmosis on the basis of a Debye model appropriately extended to handle ionic systems.

We first describe the energy barrier model as applied to the simplest case of immediate interest in cell and neurophysiology (Blaustein, 2020; Aminzare and Kay, 2024). The system contains permeant Na^+^, K^+^, and Cl^−^ions along with an impermeant anionic (negative charge) solute on one side of a membrane. The aqueous solutions are dilute, in the same range of accuracy as van’t Hoff’s equation. We then present the resulting equation for the osmotic volume flow ***J***_*v*_ (mainly due to water flow for the dilute solutions considered). For the sake of emphasis the special case of equal osmolarities is made explicit. We then exhibit the range of parameters in which the sodium pump must operate if it is to reduce the osmotic water flow ***J***_*v*_ to zero. The equation for ***J***_*v*_ has a simple appearance, but its proof, although straightforward, is not short and is presented in a separate section labeled Derivation, the theoretical equivalent of a Materials and Methods section.

## 2 Model for Donnan Osmosis in an Ionic System

We deal here with a semipermeable membrane separating two infinite aqueous compartments. The compartment on the left (see Figure 1) is called the outer compartment, and the quantities pertaining to it are designated by “out”. The compartment on the right is the inner compartment “in”. The outer compartment contains permeant Na^+^, K^+^, and Cl^−^ ions of respective concentrations (molarities) [Na^+^]_out_, [K^+^]_out_, and [Cl^−^]_out_. The inner compartment also contains these permeant ions with concentrations [Na^+^]_in_, [K^+^]_in_, and [Cl^−^]_in_, but also contains an impermeant anion Z bearing net negative charge −*zq*, where *z* is a positive number and *q* is the unit positive charge on a proton. The concentration of the impermeant anion is [Z]_in_. All outer and inner concentrations have fixed values (infinite bath model).

**Figure 1.**
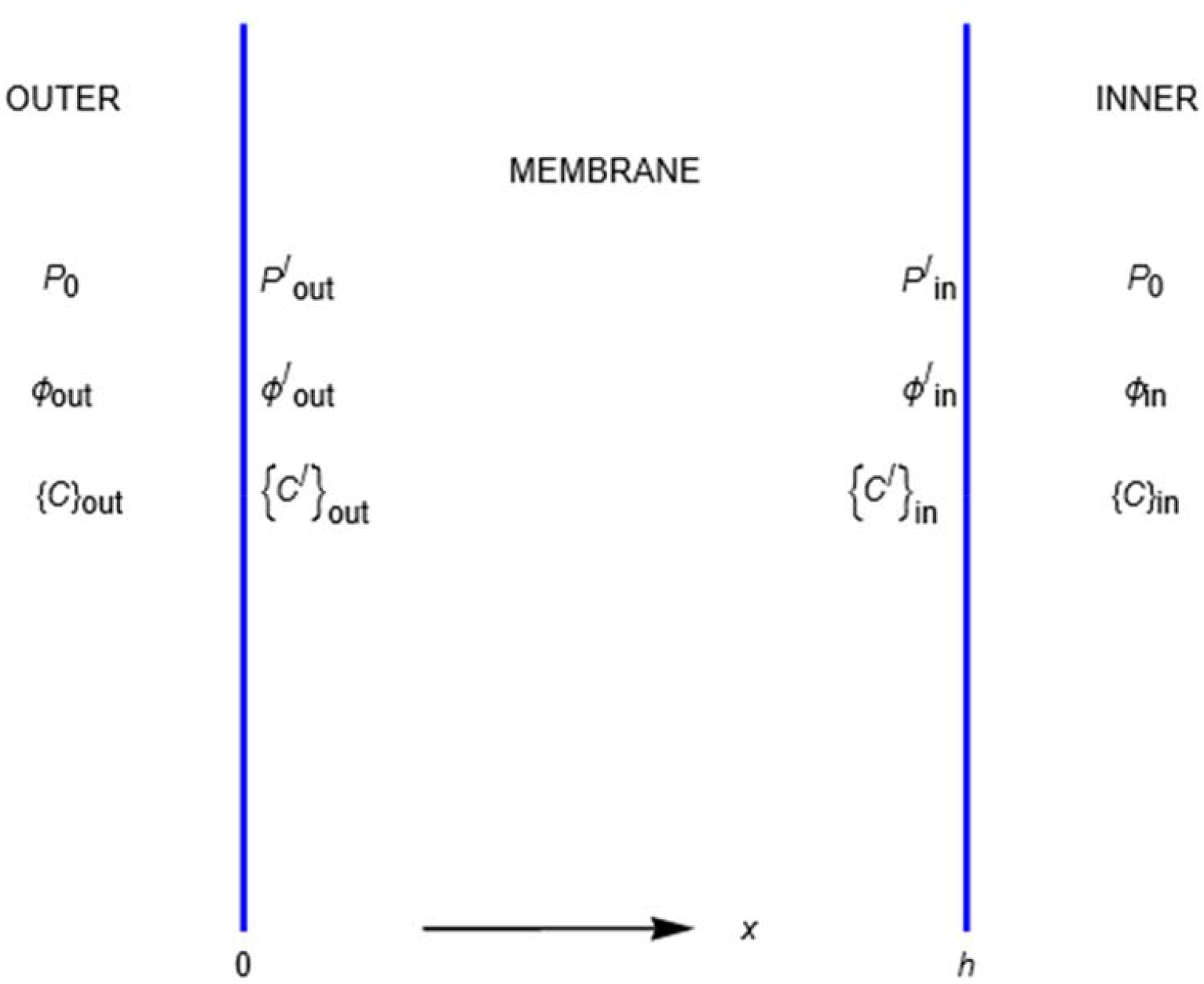
The Donnan system, showing the relevant quantities pressure, electric potential, a set of small ion concentrations {c} in the outer compartment x<0, and a set of ion concentrations in the inner compartment, x>h, which includes the concentration of impermeant anion. These quantities are discontinuous at the membrane boundaries x=0 and x=h, and their values just inside the membrane are designated by primes. The value of the impermeant anion concentration just inside the membrane is zero, but the concentrations of permeant ions inside the membrane are not zero.

The electroneutrality condition in the outer compartment is [Na^+^]_out_ + [K^+^]_out_ = [Cl^−^]_out_. In the inner compartment the electroneutrality condition is [Na^+^]_in_ + [K^+^]_in_ = *z*[Z]_in_ + [Cl^−^]_in_.

In a Donnan system at equilibrium the pressure in the inner compartment ***P***_in_ is greater than the pressure ***P***_out_ in the outer one, and the pressure difference Δ***P*** = ***P***_in_ − ***P***_out_ is called the Donnan osmotic pressure. In this paper we study the Donnan system when the osmotic pressure is not allowed to build up. Instead, the pressures in both reservoirs are maintained at equal values ***P***_0_, so that *Δ**P*** = *0*. However, just as for the simpler case of a neutral impermeant solute and no permeant ions (Manning, 1968; Manning and Kay, 2003), there are abrupt pressure drops at the membrane/solution surfaces. The pressure drops are not equal, and the resulting pressure gradient inside the membrane drives water flow.

The pressure at the surface between membrane and outer compartment is designated by 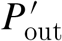, so that the pressure drop at the outer surface is 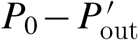. At the surface between membrane and inner compartment the pressure drops from *P*_0_ to 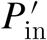. The pressure difference across the membrane from one surface to the other is then 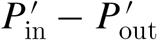, and this pressure difference is denoted by (Δ***P***)_membrane_. The expression for (Δ***P***)_membrane_ is stated as Eq. 2 below, and it is derived in a subsequent section.

The model for the membrane itself is drawn in Figure 2. It consists of an infinitely high barrier of potential energy presented to the impermeant anion Z. The infinite height prevents a Z anion from entering the space occupied by the membrane. The individual energy barrier heights for the permeant ions are arbitrarily drawn, the main point being that they are not infinitely high, allowing penetrability of the membrane space. Water molecules see no energy barrier. The potential energies represent repulsive mechanical forces between the membrane material and solute particles acting at the membrane interfaces. The relation between potential energy *u* and corresponding force *f* is *f* = −*du/dx*, where *x* is the running coordinate in Figure 2, as usual increasing from left to right. The repulsive force on a solute particle is directed from left to right at the inner membrane surface when a solute particle attempts to enter the membrane from the inner compartment. Its potential energy increases, the energy derivative *du/dx* is negative, and the force *f* is positive, pointing to the right, corresponding to repulsion. In the interior of the membrane, a solute particle is symmetrically surrounded by membrane material, so *du/dx* = *0*, and the net force *f* vanishes.

**Figure 2.**
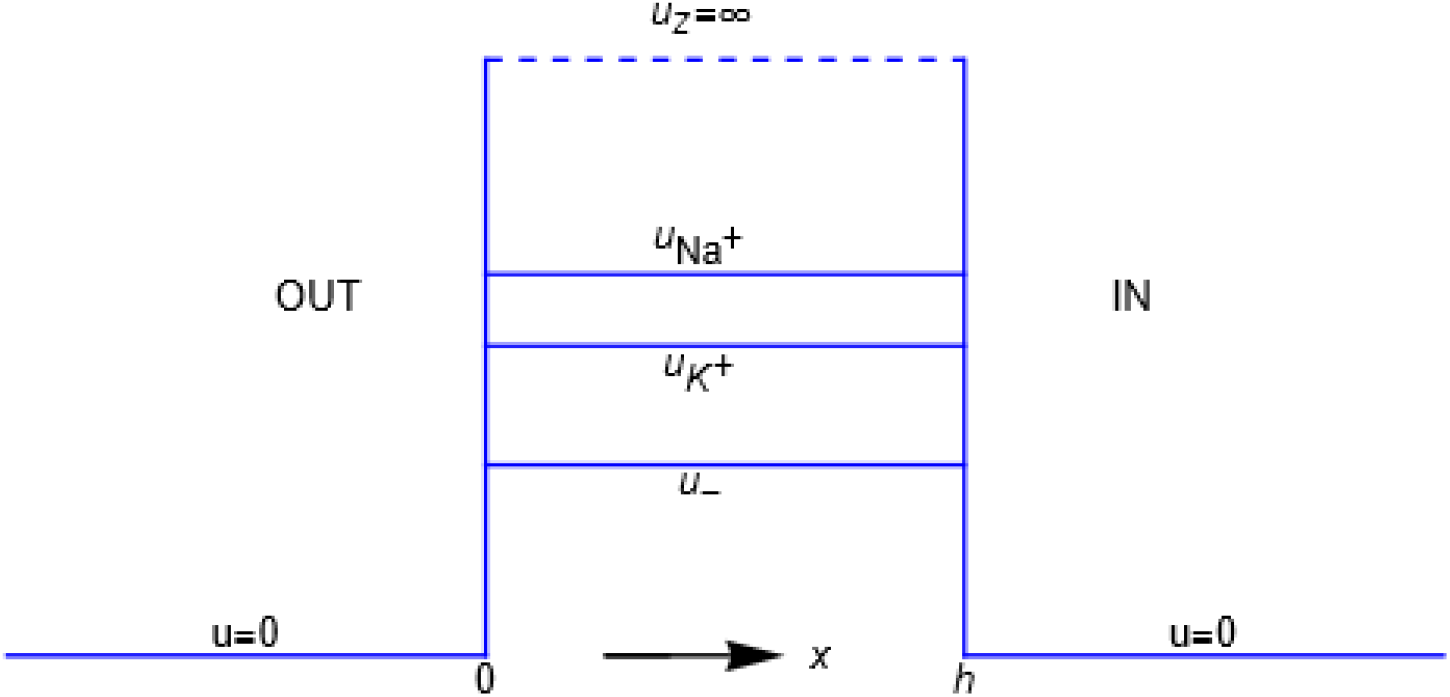
Profiles of the Debye potential energies u_*i*_ representing the mechanical interactions of the membrane with the four types of ions considered in this paper. The ordering of the potential energies of the small ions is arbitrary.

Since the mechanical forces *f* act only at the membrane/solution surfaces, they cause discontinuities there. The most obvious discontinuity is in the concentration [Z]_in_ of the impermeant anion which drops abruptly at the surface from its uniform value in the inner compartment to zero inside the membrane. The discontinuities of the membrane potential energies are evident from Figure 2. As discussed, there are also abrupt pressure drops. These discontinuities are important to understand but actually easy to handle mathematically as discussed in the section Derivation.

## 3 The Equation for Donnan Osmotic Flow

The key result from mathematical analysis of this model (see Derivation section) is an equation for the osmotic volume flow,

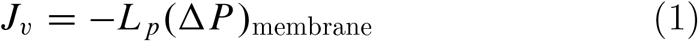

where

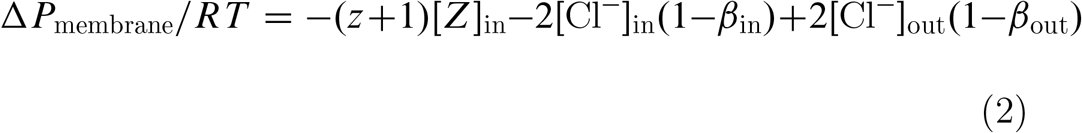

and the coefficient ***L***_*p*_ is the hydraulic permeability (***R*** is the gas constant and *T* the Kelvin temperature). It is important to understand the signs. Since the pressure drop is usually (but not always in this Donnan system) greater facing the inner compartment which contains the impermeant solute than in the outer compartment, (Δ***P***)_membrane_ is usually negative. The osmotic water flow ***J***_*v*_ is then positive, directed left to right in Figures 1 and 2 from the outer compartment into the inner compartment.

The first term on the right side of the equation for Δ***P***_membrane_ should be familiar. It is the sum of a van’t Hoff term [Z]_in_ and another van’t Hoff term *z*[Z]_in_ accounting for the *z* univalent sodium and potassium counterions required to remain in the inner compartment to neutralize (electrostatically) the charge −*zq* on the impermeant anion. Notice that it makes no sense to try to specify what proportion of the neutralizing counterions are sodium and what potassium. The neutralization is long-range electrostatic in nature, and all that can be said is that *z* among the sodium and potassium ions are rendered effectively impermeant by each mechanically impermeant Z anion.

The two further terms on the right side of the equation contain the effect of the permeant Na^+^, K^+^, and Cl^−^ ions other than those required to neutralize the charge on the Z anion. Only the concentrations of the chloride ions appear explicitly in these electrostatic terms because the combined charge of these “other” sodium and potassium ions is constrained to neutralize the charge on the chloride ions, accounting for the factor 2. The concentrations of sodium and potassium ions do appear in the definitions of the quantities *β*_in_ and *β*_out_,

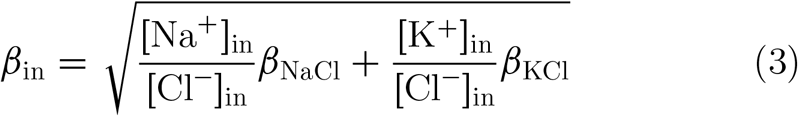

and,

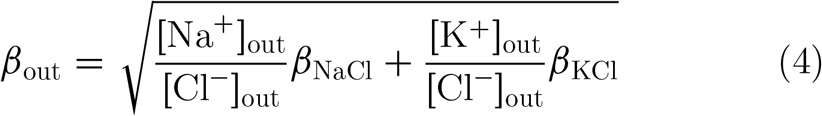

In these expressions, *β*_NaCl_ and *β*_KCl_ are partition coefficients,

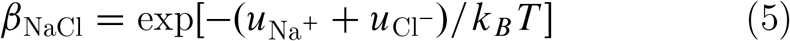

and,

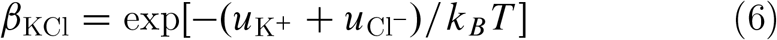

In these last two equations, *u*_Na_+, *u*_K_+, and *u*_Cl_− are the energy barrier heights for the indicated ions, and *k*_*B*_ is the Boltzmann constant (gas constant ***R*** divided by Avogadro’s number).

A useful observation follows if the membrane energy barriers are actually barriers, in other words, if all of the energies *u*_*i*_ are positive. Then the partition coefficients *β*_NaCl_ and *β*_KCl_ are less than unity, and it is easy to show from the electroneutrality conditions that the concentration-averaged partition coefficients *β*_in_ and *β*_out_ defined in Eq. 3 and Eq. 4 obey the inequalities 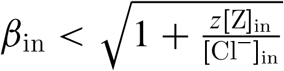 and *β*_out_ *<* 1. If all of the barrier heights *u*_*i*_ are zero, so that there are no membrane impediments to transit of the sodium, potassium, or chloride ions, the inequalities become equalities,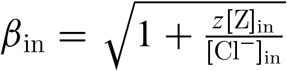 and *β*_out_ = 1..

### 3.1 Special Cases

Let us look now at two special cases, Donnan osmosis if the small ions are completely impermeant along with the impermeant anion Z, and Donnan osmosis if the small ions are freely permeant (equally as permeant as the water molecules). For impermeant sodium, potassium, and chloride ions, the corresponding Debye energy barrier levels *u* are infinite, so that *β*_in_ = *β*_out_ = *0*. The expression for volume flow reduces to,

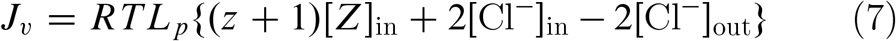

If at the opposite extreme the small ions are freely permeant, the barrier energies for them are all zero, and Donnan volume flow becomes,

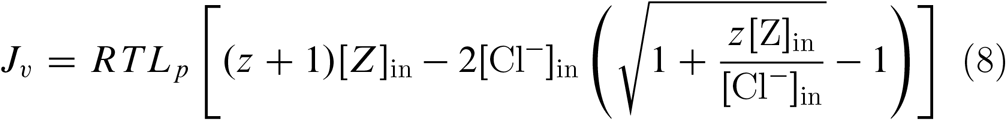

The special case Eq. 7 for impermeant small ions can be recognized as an example of the familiar van’t Hoff osmosis, or Starling’s equation for impermeant solutes. For this case, osmotic flow is indeed induced by the difference between osmolarities across the membrane, and indeed vanishes along with the osmolarity difference. On the other hand, the second term in the special case Eq. 8 for freely permeant small ions indicates a nonvanishing influence of these ions on osmotic flow even though they are freely permeant. In this case we have an effect of the Donnan electrostatic potential.

### 3.2 Numerical examples

Before setting forth on a numerical analysis of the full equation for ***J***_*v*_, it is worth making another observation about the partition coefficients *β*_NaCl_ and *β*_KCl_. They are not fully independent, since they share the energy barrier for the chloride ion. However, it may be checked that many (actually, infinitely many) combinations of positive energies *u*_Na_+, *u*_K_+, *u*_Cl_− correspond to any particular combination of positive values of *β*_NaCl_ and *β*_KCl_ less than unity (a positive value of a partition coefficient less than unity means hindered membrane permeability). In other words, every point (*β*_NaCl_, *β*_KCl_) in the square defined by *0* ≤ *β*_NaCl_, *β*_KCl_ ≤ *l* is physically accessible.

We start with an example of equal osmolarities on the two sides of the membrane. The impermeant anion Z is not necessary for the feature we wish to emphasize, so we set [Z]_in_ = *0*, [Cl^−^]_in_ = [Cl^−^]_out_ = *0*.*l* M, [Na^+^]_in_ = [K^+^]_out_ = *0*.*03* M, and [Na^+^]_out_ = [K^+^]_in_ = *0*.*07* M. The osmolarity in both inner and outer compartments is 0.2 M. In Figure 3 there is a 3D plot of Δ***P***_membrane_*/**R**T* [Cl^−^]_in_ as a function of points in a square of side unity defined by the partition coefficients *β*_NaCl_ and *β*_KCl_. At the origin *β*_NaCl_ = *β*_KCl_ = *0* and also at the point (1,1), the internal pressure gradient that drives osmotic flow (see eq 1) is indeed equal to zero. This observation is expected, since when the partition coefficients are both zero, the ions are impermeant. Zero osmotic flow is expected also if both partition coefficients have value unity, signifying freely permeant ions (same as water). It is also true that ***J***_*v*_ = *0* along the entire line *β*_NaCl_ = *β*_KCl_. But everywhere else, i.e., whenever *β*_NaCl_ ≠ *β*_KCl_, the osmotic flow does not vanish. Equal osmolarities is a false criterion for no osmotic flow.

**Figure 3.**
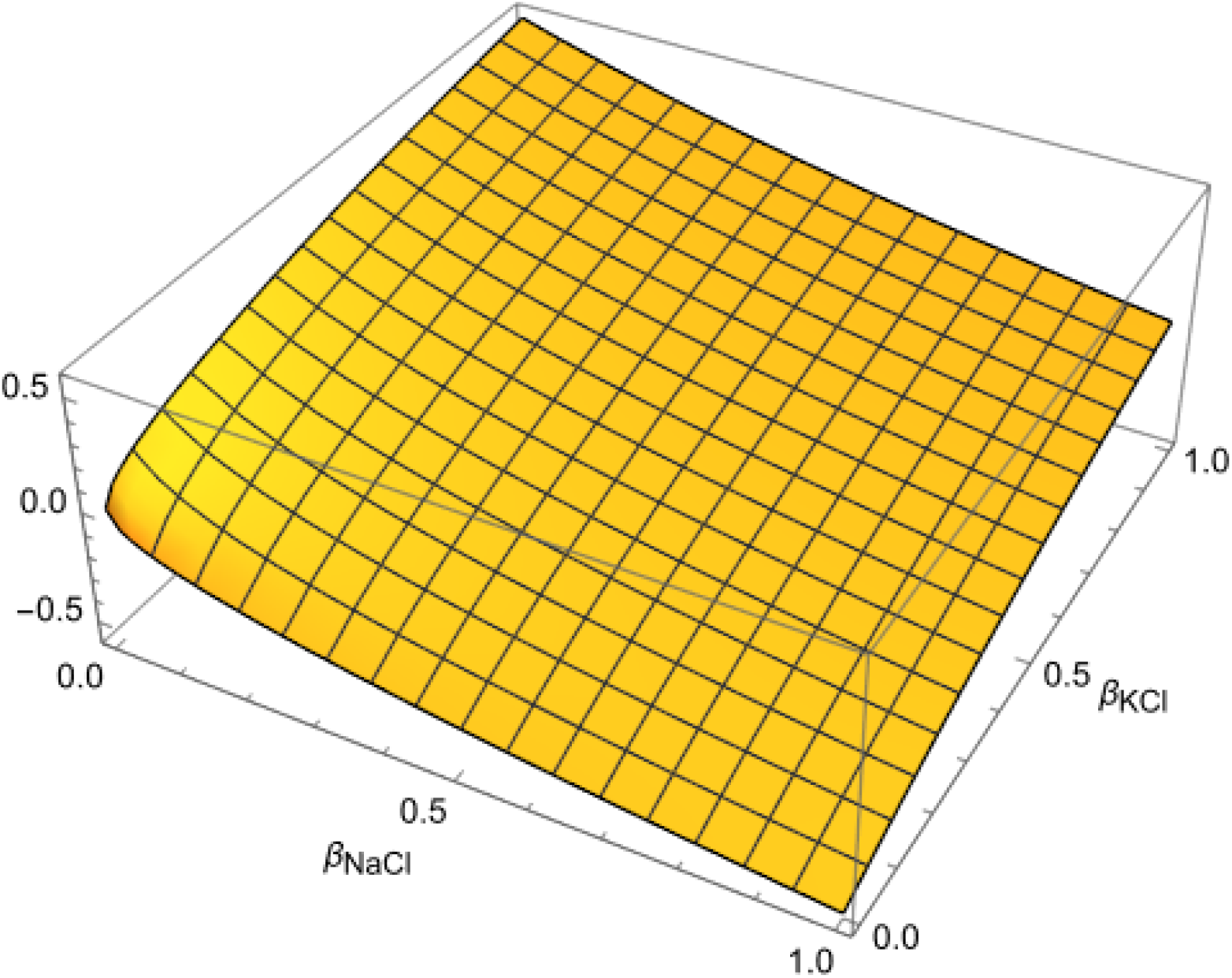
The surface is the normalized internal membrane pressure difference ΔP_membrane_ as a function of the NaCl and KCl partition coefficients for equal osmolarities in inner and outer compartments. Osmotic volume flow is proportional to −ΔP_membrane_ (see Eq. 1).

Figure 4 shows a 3D plot of Δ***P***_membrane_*/RT* [*Z*]_in_ from eq 2. The impermeant anion Z is present at concentration 0.144 M. The ion concentrations as listed by Blaustein et al. (2020) for a typical mammalian cell are [Cl^−^]_in_=0.006 M, [Cl^−^]_out_=0.106 M, [Na^+^]_in_=0.010 M, [K^+^]_in_=0.140 M, [Na^+^]_out_=0.145 M, [K^+^]_out_ = *0*.*005* M. Interestingly, the combined sodium and potassium concentrations are equal in the extracellular fluid (“out” in this paper) and the intracellular cytoplasm (“in” here), but Na^+^ is enriched outside the cell, and K^+^ inside. The concentration of impermeant anion has been chosen to meet the requirement of electroneutrality for *z*=1, a value suggested by Aminzare and Kay (2024). We see from Figure 4 that the values of Δ***P***_membrane_ are negative nearly everywhere, driving osmotic flow ***J***_*v*_ into the inner compartment containing the impermeant anion. Only along the axis *β*_NaCl_ = *0* is ***J***_*v*_ zero or very small, whereas along this axis the value of *β*_KCl_ ranges from zero (impermeant) to unity (freely permeant).

**Figure 4.**
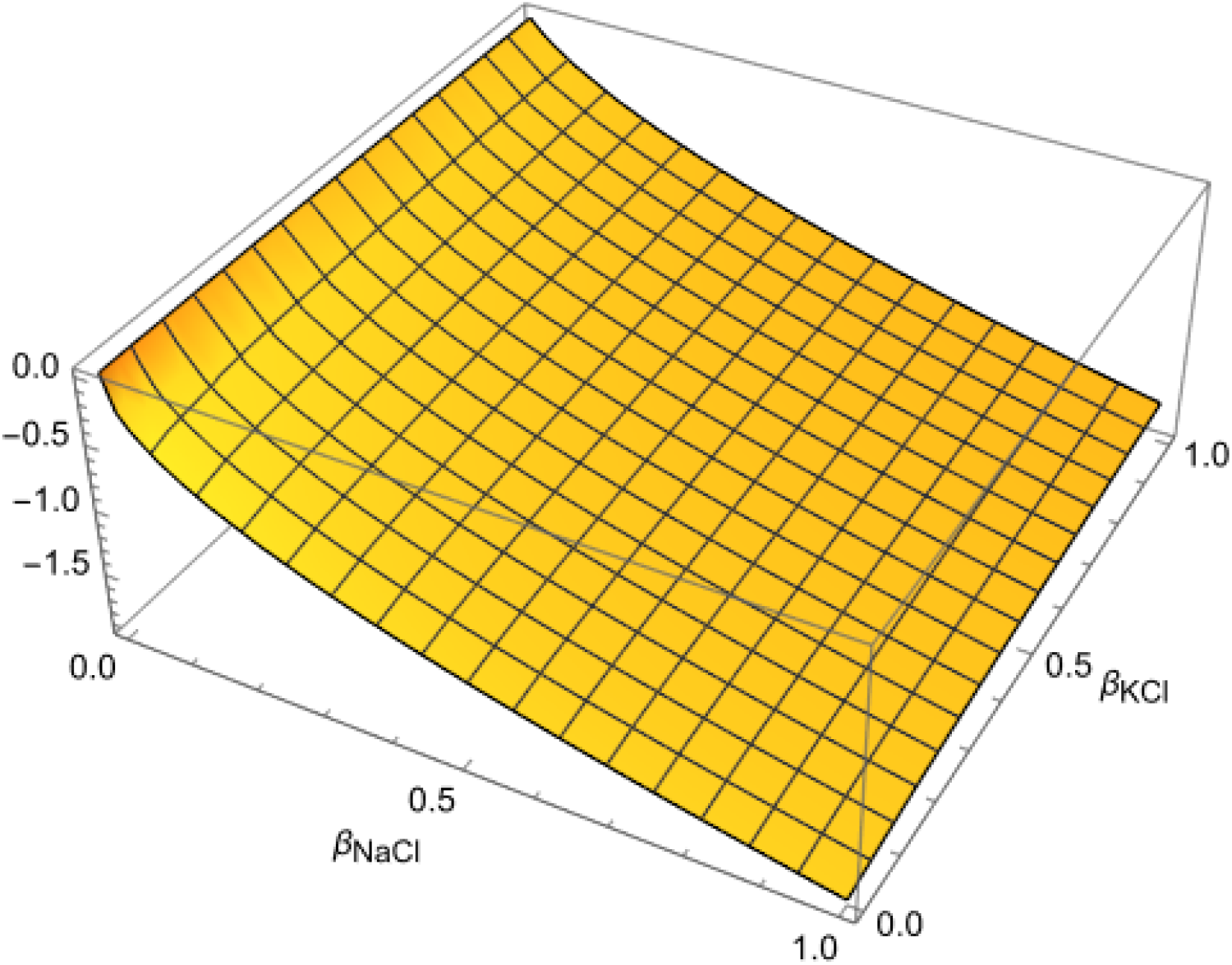
The surface plotted here is the normalized internal pressure difference ΔP_membrane_ from Eq. 2 for ionic concentrations typical of animal cells as a function of the NaCl and KCl partition coefficients. Negative values of ΔP_membrane_ signify osmotic volume flow into the cell (see Eq. 1).

The results in Figure 4 are consistent with an informal interpretation of the action of the sodium pump (Blaustein et al., 2020): “… Na^+^ ions, the most abundant permeant cations outside the cell, are extruded from the cell by active transport as rapidly as they leak in … functionally equivalent to making the cell membrane impermeable to Na^+^ ions”, and thus “forestalling osmotic catastrophe”. Indeed, the graph in Figure 4 shows that osmotic water flow into the cell (inner compartment) is reduced to zero or near-zero values when *β*_NaCl_ = *0*, that is, when the Na^+^ ion is impermeant.

As our last example, the implication of Donnan osmosis for transport across capillary walls may be of interest. Capillary walls are highly porous, and small solutes, including ions, pass through easily (Blaustein et al., 2020). The special case Eq. 8 for freely permeant ions is therefore applicable. Using the concentrations specified by Blaustein et al. in their Box 3.4, and assuming the value *z* = *l* (Aminzare and Kay, 2024), we calculate ***J***_*v*_*/RTL*_*p*_=0.82 (this flow from osmosis does not include the flow caused by the hydrostatic blood pressure). Of possibly greater interest than the number itself is that the volume flow would have been much greater (by a factor of two) if the negative contribution of the second term in Eq. 8 had been absent. Now this second term is precisely what distinguishes Donnan osmosis (in this special case) from the more familiar van’t Hoff osmosis. It accounts for the electrical effect of the permeant small ions, which is not taken into account in the usual listings of Starling forces responsible for capillary transport.

## 4 Derivation

In the derivation the notation for concentrations has been streamlined. The concentrations of the sodium and potassium ions are respectively *c*_1_ and *c*_2_, the concentration of the chloride ion is *c*_−_, and the concentration of the impermeable anion is *c*_*z*_. The units of concentration are number of molecules per unit volume, so that Boltzmann’s constant *k*_*B*_ replaces the gas constant *R*.

In the theory of the hydrodynamics of fluids, the fundamental equation for uniform volume flow in a direction *x* expresses the imbalance of forces on a volume element of fluid,

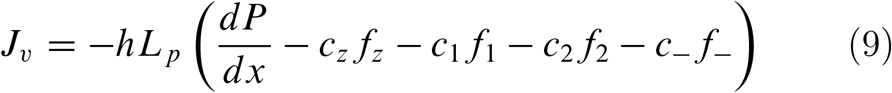

For flow across a membrane of thickness *h*, the coefficient *L*_*p*_ is the hydraulic permeability. The collection of terms in parentheses is the total force acting on a cross-sectional slab of width *dx* located at *x*. The pressure derivative is the net force exerted per unit area on the two planar surfaces of the slab. The second term is the mechanical force *f*_*z*_ from the membrane material exerted on molecules of the impermeant anion located within the slab. Of course the concentration *c*_*z*_ of the impermeant anion is zero inside the membrane, and the repelling force *f*_*z*_ acts only at the membrane interface with the inner compartment containing it (*x* = *h* in Fig. 1). The following three terms are the mechanical forces on the sodium, potassium, and chloride ions, and these forces also are nonzero only at the interfaces, because the environment of an ion inside the membrane is symmetric on both sides of it. The forces *f*_*i*_ made explicit in this equation are the mechanical ion-membrane forces. There are also electric forces on all of the ionic charges due to an electric field −*dø/dx*. But the slab is net electrically neutral, as the total charge of all the ions inside it sum to zero, so the total electric force on the slab is zero.

The beginning step is the elimination of the mechanical force *f*_*z*_ on the impermeant anion. The equation for the flux ***J***_*z*_ of this ion is,

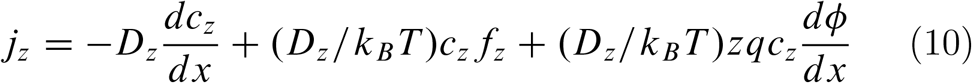

where *D*_*z*_ is a diffusion constant. But ***J***_*z*_ = *0* since the Z ion is impermeant, and the resulting equation can be solved for *c*_*z*_*f*_*z*_, which can then be substituted back into Eq. 9 to give,

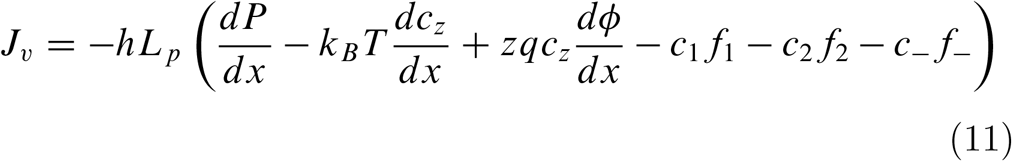

Next, we invoke electroneutrality, *zc*_*z*_ = *c*_1_ + *c*_2_ − *c*_−_, in the electric field term,

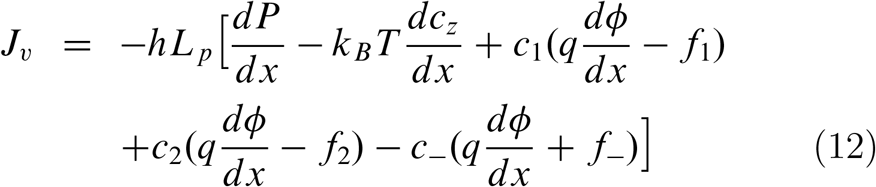

The electrical and mechanical forces on the sodium, potassium, and chloride ions can be eliminated if we look at the equations for their fluxes, which do not vanish since these ions are permeant. For example,

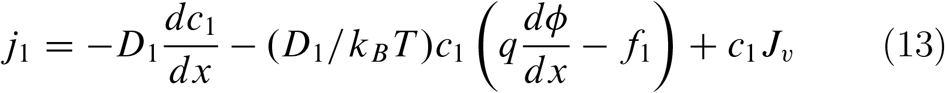

where a solvent drag term has been added (it does not affect the final result). This equation can be solved for the combined electrical and mechanical force term for the sodium ion, which is then substituted into Eq. 12. After analogous substitutions for the potassium and chloride ions, Eq. 12 becomes,

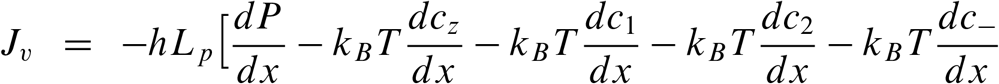

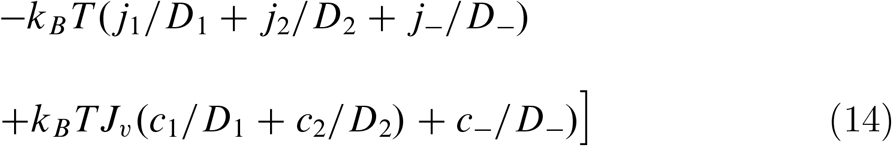

Both sides of this equation can be integrated across the surface separating the outer compartment from the interior of the membrane. The terms involving ***J***_*v*_ and the ***J***_*i*_ integrate to zero, because the integration is over an infinitesimal interval (just outside the membrane to just inside). The integral of *dP/dx* equals the difference 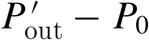, that is, the pressure drop at the outer surface. The integral of the *dc*_*z*_*/dx* term vanishes, because *c*_*z*_ is zero both in the outer compartment and inside the membrane. As for the permeant ion terms, as an example, the integral of the derivative *dc*_1_*/dx* across the outer interface equals the difference 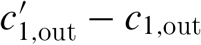, where the first term is the concentration just inside the interface, and the second is the concentration just outside. The result of the integration is,

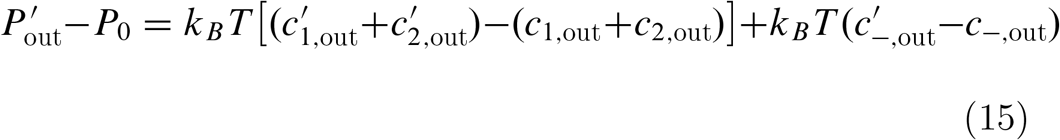

or, from electroneutrality *c*_1_ + *c*_2_ = *c*_−_ for both the primed and unprimed concentrations,

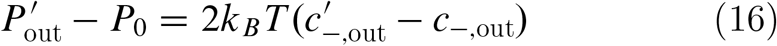

Performing an analogous integration of Eq. 14 from left to right across the inner membrane surface, with electroneutrality here reading *c*_1_ + *c*_2_ = *zc*_*z*_ + *c*_−_ outside the membrane in the inner compartment, and 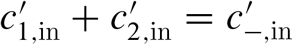 just inside the membrane at the inner surface, we get,

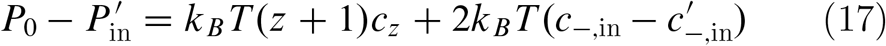

The pressure gradient inside the membrane, from one side to the other, is *(ΔP)*_membrane_, equal to 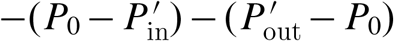, so adding the negatives of the previous two equations,

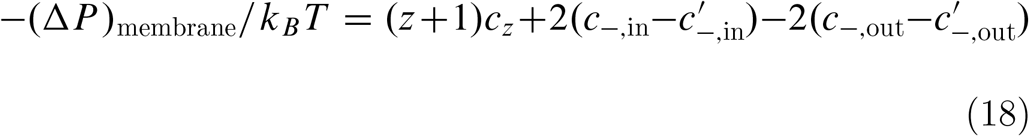

The primed chloride concentrations in this equation for *(ΔP)*_membrane_ are the concentrations just inside the membrane at the interfaces. They are not known at this point in the derivation, and the next task is to find them. We write the flux equations for the sodium and chloride ions, and add them to eliminate the electric field terms,

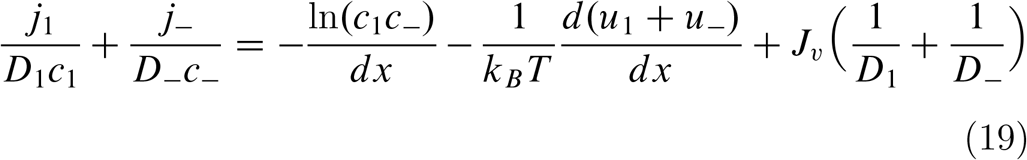

In this equation we have used *f*_1_ = −*du*_1_*/dx* and *f*_−_ = −*du*_−_*/dx* Similarly,

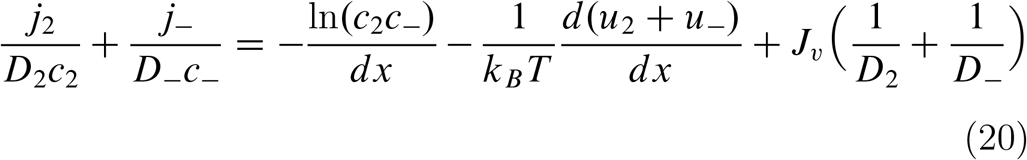

Now integrate both sides of Eq. 19 across the outer membrane surface, noting that *u*_1_ = *u*_−_ = *0* in the outer compartment, and, as above, that integration of the flow terms vanish. After exponentiating,

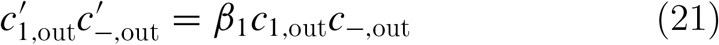

where *β*_1_ is the same partition coefficient as *β*_NaCl_ in Eq. 5 in the text. Analogous handling of the potassium and chloride ion fluxes yields,

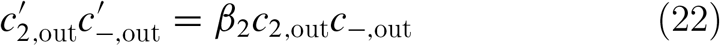

where *β*_2_ is the same partition coefficient as *β*_KCl_ in Eq. 6 in the text. Together with electroneutrality, 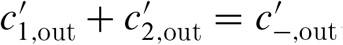 the preceding two equations give us three equations for the three unknowns 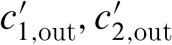 and 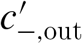, namely the sodium, potassium, and chloride ion concentrations just inside the membrane at its outer surface. Solving,

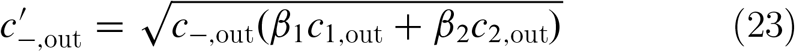

There are expressions for the sodium and potassium concentrations also, but they are not needed here. Having noticed that the electroneutrality condition inside the membrane at the inner surface has the same form as for the outer since *c*_*z*_ = *0* inside the membrane, we handle the inner surface analogously,

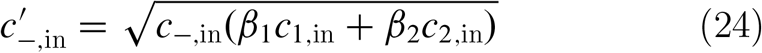

Substitution of these two equations into Eq. 18 results in Eq. 2 of the text and completes the derivation.

## 5 Discussion

It is well known in physical chemistry that ionic solutions behave very differently from solutions with only neutral solutes (Mayer, 1950). Coulomb electrostatic forces are both long-range and unusually strong. Ions are ubiquitous in biological systems, and it is presumably important to understand the consequences of their electrical charges in areas other than neuroscience. If present, ionic interactions are a dominant influence in osmotic flow. Osmosis in a Donnan ionic system is driven by an abrupt pressure drop at the membrane interface, just as is classic osmosis in systems with neutral solutes (Manning and Kay, 2023). However, the equation presented here for Donnan osmosis in a system containing permeant Na^+^, K^+^, and Cl^−^ ions along with an impermeant anion is quite different from the expression for volume flow stemming from differences in neutral solute concentrations (Kedem and Katchalsky, 1958; Manning, 1968).

We have seen that even if the small ions are freely permeant, like the solvent water itself, their electrostatic charges interact with the Donnan potential to contribute to the osmotic flow. Further, equal osmolarities on both sides of the membrane do not imply vanishing osmotic flow. As an application of Donnan osmosis, we showed that given the presence of ionic concentrations typical of mammalian cells, osmotic volume flow into the compartment containing the impermeant anion can be prevented if the membrane is provided with a pump that effectively renders the Na^+^ ions impermeant. We also showed that the contribution of Donnan charge effects to osmotic flow across capillary walls is very large.

## Appendix

It is remarkably difficult to find a complete treatment of Donnan equilibrium (but see Overbeek, 1956). Expressions for the three equilibrium Donnan effects in ideal dilute conditions are listed here from my unpublished lecture notes. The Donnan system consists of a semipermeable membrane separating an aqueous salt solution like NaCl in an “outer” compartment from a corresponding solution in an “inner” compartment that also contains an impermeant anion Z with charge −*zq, z* a positive number, *q* the charge on a proton, and concentration *c*_*z*_. The sodium and chloride ions are permeant. Equilibrium has been established between inner and outer compartments. Concentrations in the outer compartment are marked with a “prime”. Concentrations in the inner compartment are unmarked. The salt concentration in the outer compartment is 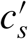 (equal to the sodium and chloride ion concentrations there), assumed known. The salt concentration in the inner compartment is *c*_*s*_ (equal to the chloride concentration there, the sodium concentration being equal to *zc*_*z*_ + *c*_*s*_).

Define 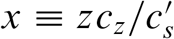 (note the different meaning of *x* here than in the text). Define 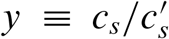 as the Donnan salt exclusion ratio (Donnan ion distribution). Then at equilibrium, *y(x)* is the function of *x* given by the solution of the quadratic equation (*x* + *y*)*y* = 1.

The Donnan electrostatic potential difference *ø* − *ø*′ between inner and outer compartments is given by,

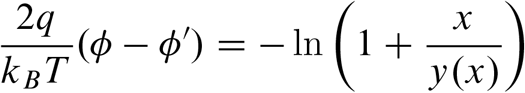

(*q/k*_*B*_*T* = *F/RT*, where *F* is the Faraday).

The Donnan osmotic pressure is given by,

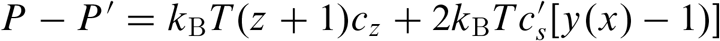

## Acknowledgement

The author is grateful to Alan R. Kay for numerous stimulating discussions.

## References

Aminzare, Z., and A.R. Kay. 2024. Mathematical modeling of intracellular osmolarity and cell volume stabilization: The Donnan effect and ion transport. J. Gen. Physiol. Vol. 155 No. 10 e202313332.

Blaustein, M.P., J.P.Y. Kao, and D.R. Matteson. 2020. Cellular Physiology and Neurophysiology. Third edition. Elsevier, St. Louis.

Debye P. 1923a. Kinetische theorie der gesetze des osmotischen drucks bei starken elektrolyten. Phys. Z. 24:334338.

Debye, P. 1923b. Theorie cinetique des lois de la pression osmotique des electrolytes forts. Recl. Trav. Chim. Pays Bas. 42:597604. 10.1002/recl.19230420711

Donnan, F.G. 1995. Theory of membrane equilibria and membrane potentials in the presence of non-dialysing electrolytes. A contribution to physical-chemical physiology. J. Membrane Sci. 100:45–55 (This paper is an English language translation from the German of Donnan’s original 1911 paper, cited therein).

Feher, J. (2018). Quantitative Human Physiology. Second Edition. Academic Press, Cambridge, MA.

Kedem, O., and A. Katchalsky. 1958. Thermodynamic analysis of the permeability of biological membranes to non-electrolytes. Biochim. Biophys. Acta 27:229–246.

Keener, J., and J. Sneyd. 2009. Mathematical Physiology I. Cellular Physiology. Second edition. Springer, NY.

Manning, G.S. 1968. Binary diffusion and bulk flow through a potential energy profile: A kinetic basis for the thermodynamic equations of flow through membranes. J. Chem. Phys. 49:2668–2675.

Manning, G.S., and A.R. Kay. 2023. The physical basis of osmosis. J. Gen. Physiol. Vol. 155 No. 10 e202313332.

Mayer, J.E. (1950). The theory of ionic solutions. J. Chem. Phys. 18:1426–1436.

Milo, R., and R. Phillips. 2019. Cell Biology by the Numbers. Taylor & Francis, UK.

Nagasawa, M., A. Takahashi, M. Izumi, and I. Kagawa. 1959. Colligative properties of polyelectrolyte solutions. VI. Donnan membrane equilibrium. J. Polymer Sci. 38:213–228.

Overbeek, J.Th.G. 1956. The Donnan Equilibrium. Prog. Biophys. Biophys. Chem. Chapter 3, pp. 57–84.

Patterson, G. 2007. Physical Chemistry of Macromolecules. CRC Press, Boca Raton.

Scatchard, G. 1946. Physical chemistry of protein solutions. I. Derivation of the equations for the osmotic pressure. J. Am. Chem. Soc. 68:2315–2319.

Tanford, C. 1961. Physical Chemistry of Macromolecules. Wiley, NY.

Weiss, T.R. 1996. Cellular Biophysics. Volume 1: Transport. MIT Press, Cambridge, MA

